# A divergent hepatitis D-like agent in birds

**DOI:** 10.1101/423707

**Authors:** Michelle Wille, Hans J. Netter, Margaret Littlejohn, Lilly Yuen, Mang Shi, John-Sebastian Eden, Marcel Klaassen, Edward C. Holmes, Aeron C. Hurt

## Abstract

Hepatitis delta virus (HDV) is currently only found in humans, and is a satellite virus that depends on hepatitis B virus (HBV) envelope proteins for assembly, release and entry. Using meta-transcriptomics, we identified the genome of a novel HDV-like agent in ducks. Sequence analysis revealed secondary structures that were shared with HDV, including self-complementarity and ribozyme features. The predicted viral protein shares 32% amino acid similarity to the small delta antigen of HDV and comprises a divergent phylogenetic lineage. The discovery of an avian HDV-like agent has important implications for the understanding of the origins of HDV and subviral agents.

**Importance:** Hepatitis delta virus (HDV) is currently only found in humans, and coinfections of HDV and Hepatitis B virus (HBV) in humans result in severe liver disease. There are a number of hypotheses for the origin of HDV, although a key component of all is that HDV only exists in humans. Here, we describe a novel deltavirus-like agent identified in wild birds. Although this agent is genetically divergent, it exhibits important similarities to HDV, such as the presence of ribosymes and self-complementarity. The discovery of an avian HDV-like agent challenges our understanding of both the origin and the co-evolutionary relationships of subviral agents with helper viruses.

## Introduction

Hepatitis delta virus (HDV) is a human-specific pathogen and the sole member of the genus *Deltavirus*. The HDV genome is unique among known animal viruses but shares similarities with plant subviral pathogens named viroids (1, 2). The single stranded, circular RNA genome of HDV is approximately 1700 nucleotides and is therefore the smallest virus infecting mammals. It is ~70% self-complimentary, and forms a highly base paired rod-like structure. It encodes two proteins (small and large delta antigen, S-HDAg and L-HDAg, respectively) from a single open reading frame. HDV is regarded as a subviral pathogen that requires the envelope proteins from the helper hepatitis B virus (HBV) for assembly and release, and subsequently for entry into the host cell (2).

In humans, coinfection with HDV and HBV causes more severe liver disease than is seen in individuals infected with HBV alone, and 15-20 million individuals are estimated to be co-infected with both viruses (3, 4). There are currently 8 broad clades, or genotypes, of HDV, with the greatest diversity found in Africa (5). However, the evolution of this virus is poorly described, and its origin largely unknown (6). Current hypotheses for the origin of HDV rely on host restriction, that is, that this virus is only found in humans and include: viroid-like RNA having captured host signalling mRNAs (7), direct origination from the human transcriptome (8), or evolution from a circular host RNA found in (1, 9). Given the dependence of HDV on HBV for replication, there has likely been co-evolution between HDV and the helper HBV, although the nature of this relationship has been largely unexplored.

In this study we describe a divergent HDV-like agent found in wild birds and in the absence of duck HBV. We demonstrate a number of featured shared between HDV and this avian HDV-like agent. This finding has important ramifications for our understanding of both the origin and the co-evolutionary relationships of subviral agents and helper viruses.

## Results and Discussion

As part of an avian meta-transcriptomic study we identified a genome related to that of HDV, indicating that a novel and divergent HDV-like agent is present in the bird population. RNA sequencing of the rRNA depleted library resulted in 20,945,917 paired reads, which were assembled into 163,473 contigs, 279 of which were most similar to virus sequences in the GenBank non-redundant protein database (nr). Avian viral transcripts were highly abundant in the library, largely representing influenza A virus. RNA from an avian HDV-like agent was ten-fold less abundant than that of influenza A virus, but was more abundant than RNA from the host reference gene RPS13 that is stably expressed in ducks (10) (Fig 1B). Exogenous duck hepatitis B virus (DHBV) was not identified in this meta-transcriptomic library. The full genome of the novel avian HDV-like agent was represented by a single contig (GenBank accession MH824555), wherein the virus genome was duplicated, the result of sequencing a circular genome of 1706 nucleotides (Fig 1A). Unlike HDV with a GC content of around 60% (11), the GC content of the avian HDV-like agent is only 51%, with no significant peaks or troughs of GC content anywhere in the genome (Fig 1A). A conserved domain search identified the ORF in the virus genome as representing the HDAg (e-value 3.40×10^−10^). The predicted avian HDV-like protein (avHDAg) shares 32.2% amino acid similarity to characterized HDAg proteins in the Genbank nr database. This phylogenetic analysis indicated that avHDAg was highly divergent from all HDAgs encoded by known human HDV genotypes (Fig 1C).

**Figure 1.**
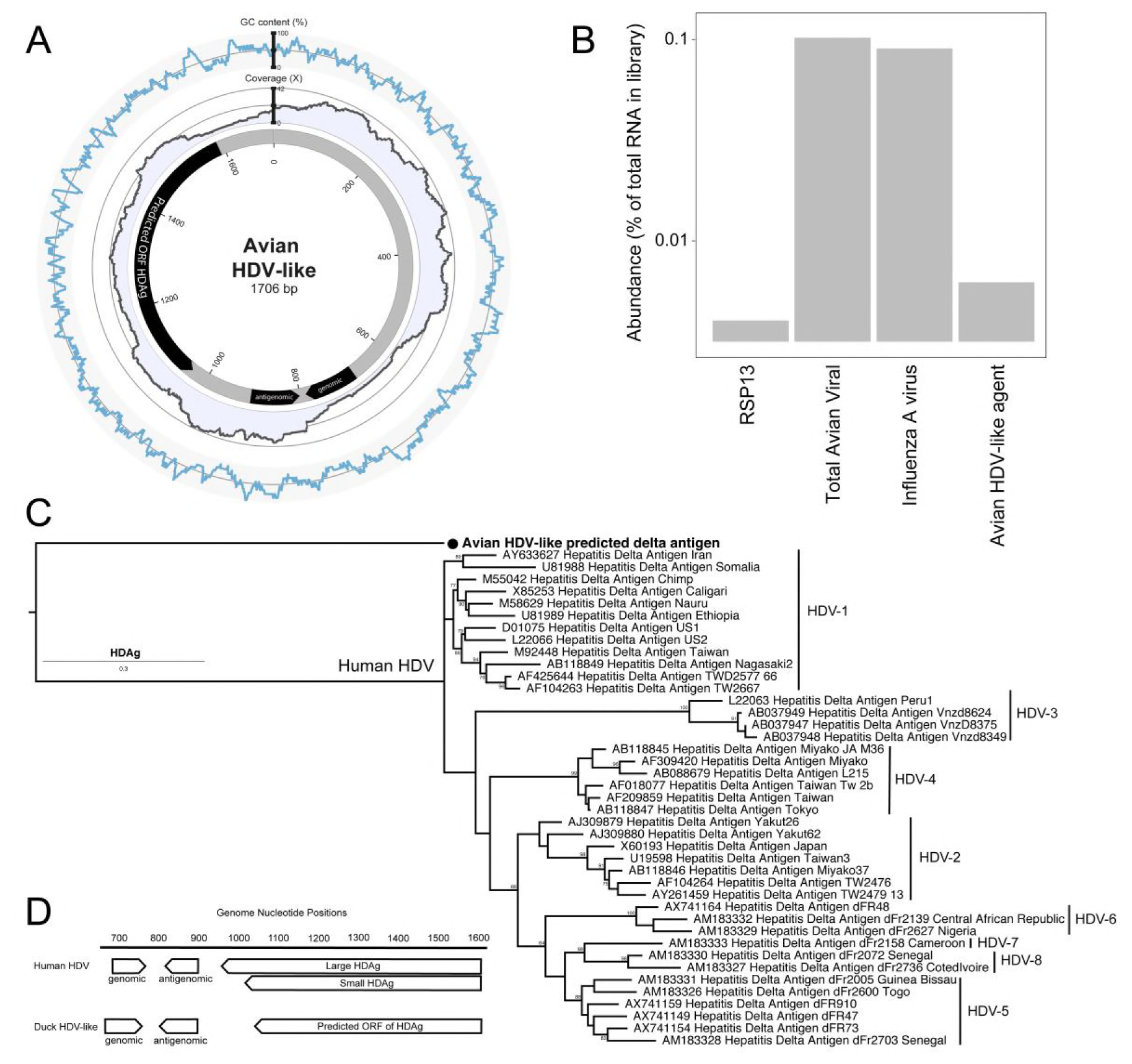
Characteristics of the genome of an avian HDV-like agent. (A) Avian HDV-like agent genome, annotated with ORFs and genomic and antigenomic ribozyme sites. Metadata rings include the read coverage, followed by GC content. (B) Abundance of transcripts in the metatranscriptomic library. Total avian viral abundance was dominated by that of influenza A virus. However, the abundance of HDV is higher than that of Ribosomal protein S13 (RPS13), a stably expressed reference gene in Mallards (*Anasplatyrhynchos*). (C) Maximum likelihood phylogeny of the HDAg protein. Representative human HDAg sequences included fall into the currently described clades HDV1 - 8 (5). The scale bar represents the number of amino acid substitutions per site. The phylogeny is rooted between the human and avian viruses. (D) Location of genomic and antigenomic ribozyme sequences, and the predicted ORF of the delta antigen in the avian HDV-like genome, compared to their location in the HDV genome sequence (GenBank accession X04451.1).

Despite this divergence, the avian HDV-like agent and HDV shared many features. In accordance with the unbranched, rod-like genome structure described for HDV, we demonstrate that the predicted that the circular RNA genome of the HDV-like agent also folded into a classic unbranched rod-like structure (Fig. 2). Importantly, consistent with HDV, the genome of the avian HDV-like agent had the capacity to express a protein, avHDAg, and contained sequences reminiscent of the HDV genomic and antigenomic ribozymes (12) and HDV-like ribozymes (13). To be consistent with the HDV nomenclature, we regard the sequence with the avHDAg ORF as the antigenome. Given that the antigenomic HDV ribozyme is located approximately 100 nucleotides downstream of the small HDAg ORF in the 3’-direction, we examined the corresponding antigenomic region of the avian HDV-like agent for the presence of ribozyme like sequences (avHDAg ORF between nucleotides 1033 and 1590, Fig 1A). Two segments in the avian HDV-like genome were identified as potential genomic and antigenomic ribozyme sequences. Their sizes and locations are very similar to the ribozymes in the reference HDV genome sequence (GenBank accession X04451.1) (Fig 1D, Fig 3). The potential genomic and antigenomic ribozyme sequences were approximately 88 and 95 nt in length, respectively (compared to 85 and 89 nt of the human HDV ribozyme sequences (12)), and the calculated free energies were -40.53 kcal/mol and -38.68 kcal/mol, respectively. When the inferred structures were re-drawn based on the canonical secondary structures of the human HDV ribozymes (13), both potential genomic and antigenomic duck ribozyme sequences displayed the ability to be folded into classic HDV ribozyme secondary structures (Fig 3). This includes the five paired (P) segments forming two coaxial stacks (P1 stacks on P1.1 and P4, while P2 stacks on P3), with these two stacks linked by single-stranded joining (J) strands J1/2 and J4/2, as described by (13).

**Figure 2.**
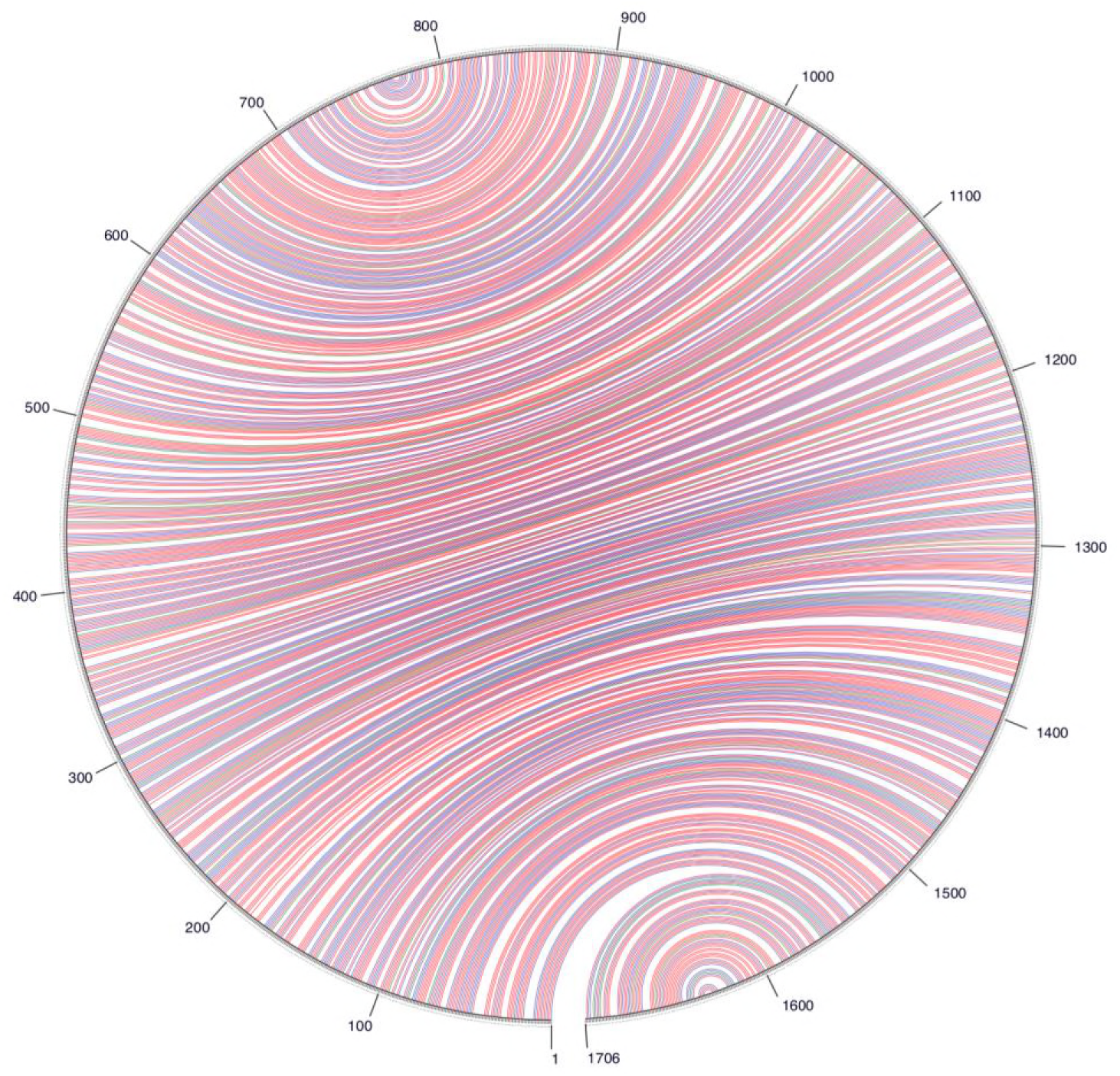
A circle graph showing the base pairing of the circular RNA genome structure of the avian HDV-like agent into an unbranched rod-like structure. The circle circumference represents the genome sequence, and the arcs represent the base pairing. Colouring of arcs are: red for G-C pairing, blue for A-U pairing, green for G-U pairing, and yellow for other type of pairings.

**Figure 3.**
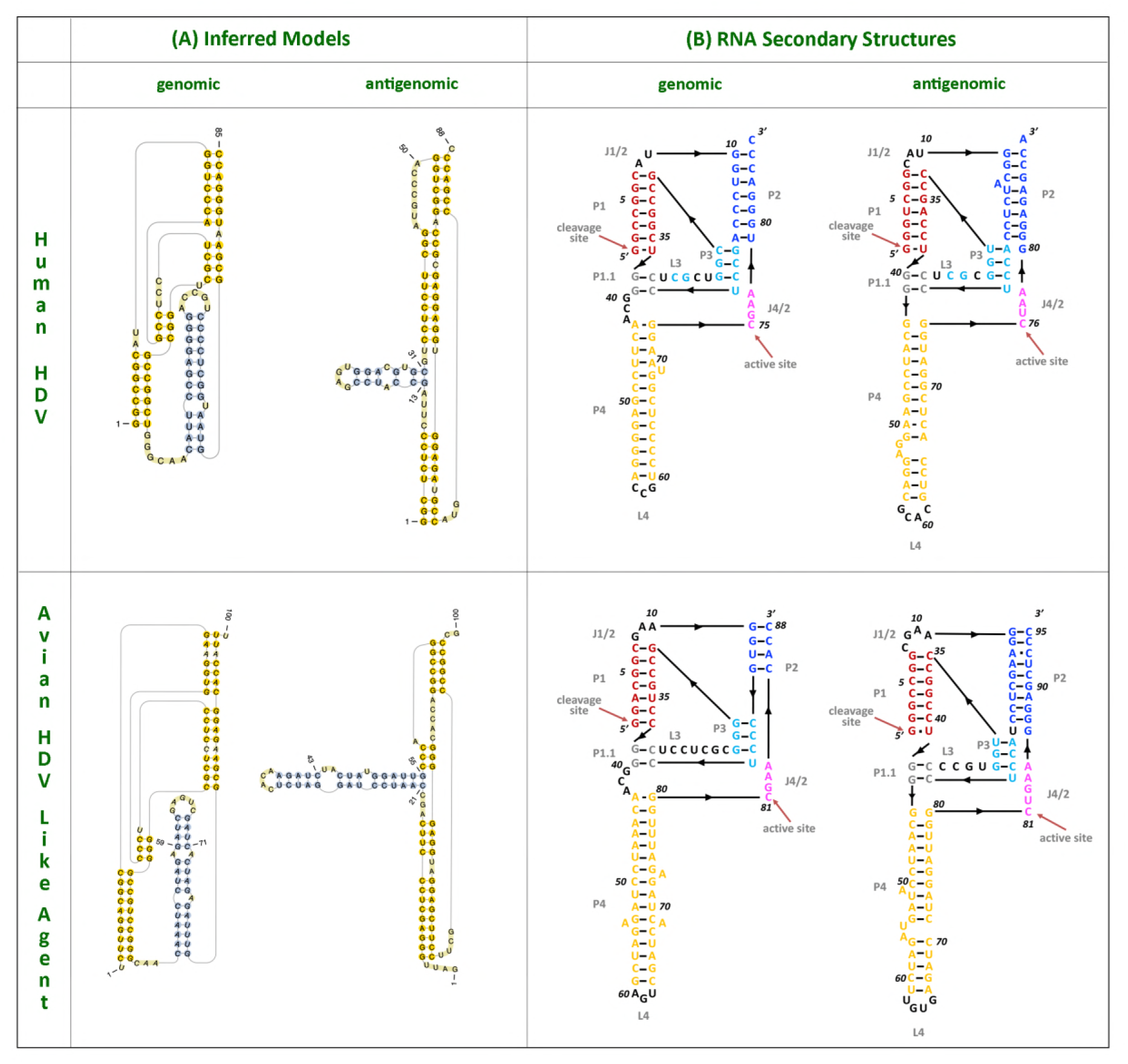
HDV ribozymes. (A) Secondary structures of the genomic and antigenomic ribozymes inferred using the TT2NE algorithm. The HDV ribozyme models were used as reference to screen for the ribozyme sequences in the avian HDV-like genome sequence. (B) Re-drawn secondary structures of the genomic and antigenomic ribozymes based on the secondary structures shown in review by Webb and Luptak (13).

The avian HDV-like genome contains a predicted ORF for the avHDAg which encodes 185 amino acids. The start AUG is located within a Kozak consensus sequence with a G in the +4 position, and Adenine in position -3 (14). Downstream of the coding sequence after 105 nucleotides, the RNA genome encodes signals which are typical for the 3’-end of eukaryotic mRNAs required for adding the poly(A) tail, the highly conserved 5’-AAUAAA recognition sequence followed by the 5’-CA-3’ cleavage sequence after 14 nucleotides (15, 16). In contrast to HDV, the genome of the avian HDV-like agent does not contain an editing site as identified for HDV. For HDV, the editing event converts the ‘UAG’ stop codon into the tryptophan ‘UGG” codon (17), which extends the reading frame by an additional 18 amino acids, resulting in the synthesis of the L-HDAg. However, the avian HDV-like genome potentially provides additional reading frames by +1 and +2 frame shifts extending the ORF of 185 amino acids by 18 and 65 amino acids, respectively. The nucleotide sequence encoding the additional 65 amino acids overlaps with the poly (A) signal sequence, and is therefore less likely to be translated from a functional mRNA molecule. The C-terminal region of the 185 amino acid protein, including the potential frame-shifted extensions do not contain a C-X-X-X isoprenylation site, which is required for the synthesis of a functional L-HDAg to support HDV assembly and release (18). Consistent with the presence of a coiled-coil domain in the N-terminal region of the HDAg, the protein avHDAg also contains a coiled-coil domain predicted in the N-terminal region between amino acids 22 and 44, probably facilitating dimerization (Fig 4) (19, 20).

**Figure 4.**
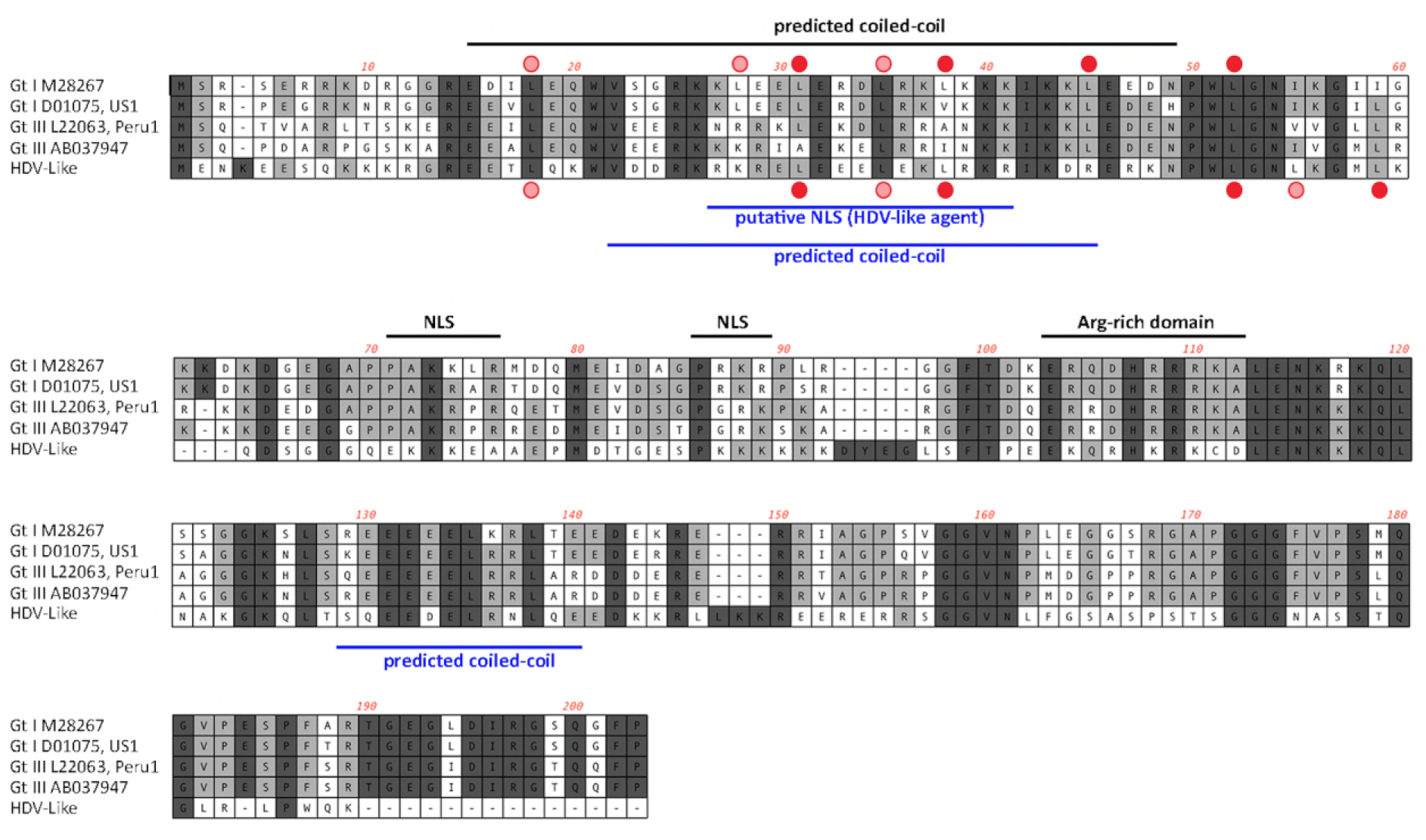
Alignment of the amino acid sequences (small delta antigen) translated from the genomes of HDV and the avian HDV-like agent. The potential coiled coil region is highlighted, including the presence of leucines in a correct spacing for a leucine zipper (filled red circle). The delta antigen does not have a strict requirement for leucine in the d position of the heptad repeat. Additional leucines shown by circles in light red. NLS: Nuclear localisation signal.

There are a number of hypotheses for the origin of HDV, including viroid-like RNA captured host signalling mRNAs (7), that HDV originated directly from the human transcriptome (8), or evolved from a circular host RNA found in hepatocytes that was able to replicate (1, 9). A central component of these hypotheses is that HDV exists only in humans. However, the discovery of an HDV-like genome in birds, with distinct similarities to the HDV genome, such as self-complementarity and ribozyme folding, but also clear differences (no ORF extension in the same frame downstream of the stop codon) suggests a divergent evolutionary pathway of HDV and HDV-like pathogens. As we were not able to detect DHBV genomes in the avian metatranscriptomic library, the identification of the genome of the avian HDV-like agent in oropharangeal and cloacal samples may indicate that the avian HDV-like agent does not depend on a hepadnavirus for the completion of its replication cycle. As such, the discovery of the genome of an avian HDV-like agent has important implications for our understanding of both the origin and the co-evolutionary relationships of the subviral agents with helper viruses, including the dependence of HDV on the HBV envelope protein.

## Materials and Methods

### Ethics statement

This research was conducted under approval of Deakin University Animal Ethics Committee (permit numbers A113-2010 and B37-2013). Banding was performed under Australian Bird Banding Scheme permit (banding authority numbers 2915 and 2703). Research permits were approved by Department of Environment, Land, Water and Planning Victoria (permit numbers 10006663 and 10005726).

### Sample selection, RNA library construction and sequencing

Waterfowl were captured at the Melbourne Water Western Treatment Plant, Victoria, Australia, in 2012-13. Oropharangeal and cloacal samples were collected from Grey Teal (*Anas gracilis*), Chestnut Teal (*A*. *castanea*) and Pacific Black Ducks (*A. superciliosa*) with no signs of disease. RNA was extracted using the MagMax *mir*Vana Total RNA isolation Kit (Thermo Scientific), assessed for RNA quality, and 10 samples with the highest concentration were pooled using equal concentrations using the RNeasy MinElute Cleanup Kit (Qiagen). Libraries were constructed and sequenced as per Shi *et al*. 2016 (21). Reads have been deposited in the Short Read Archive BioProject PRJNA472212.

### RNA virus discovery

Contigs were assembled, identified, and abundance calculated as per Shi *et al*. 2016 (21). Briefly, sequence reads were demultiplexed and trimmed with Trimmomatic followed by *de novo* assembly using Trinity (22). No filtering of host/bacterial reads was performed before assembly. All assembled contigs were compared to the entire non-redundant nucleotide (nt) and protein (nr) database with using blastn and Diamond (23), respectively, setting an e-value threshold of 1×10^−10^ to remove potential false positives. Abundance estimates for all contigs were determined using the RSEM algorithm implemented in Trinity. All contigs that returned blast hits with paired abundance estimates were filtered to remove all bacterial and host sequences. The virus list was further filtered to remove viruses with invertebrate (21), plant or bacterial host association using the Virus-Host database (http://www.genome.ip/virushostdb/).

To compare viral abundance to that of the host, a blast database was created containing Ribosomal Protein S13 (RPS13) from both Mallard (taxid: 8839) and Chicken (*Gallus gallus*) (taxid: 9031), which has found to be stably expressed in the Mallard (*Anas playrhynchos*) lower gastrointestinal tract (10).

### Characterization of novel Hepatitis D-like virus

Contigs greater than 1000bp in length were inspected (Geneious R10). Virus reads were mapped back to the HDV-like contig using the Geneious mapping function to corroborate the contig sequence and to calculate read coverage. Open reading frames were predicted within Geneious, and interrogated using the conserved domain database (CDD, https://www.ncbi.nlm.nih.gov/Structure/cdd/wrpsb.cgi), with an expected value threshold of 1×10^−5^ Reference sequences of Hepatitis D representing all eight major clades were downloaded from GenBank (Table S1). The translation of the HDAg proteins were aligned using MAFFT (24), and gaps trimmed using trimAL (removing gaps that occur in more than 20% of sequences or with a similarity scores lower than 0.005, unless this removes more than 40% of columns) (25). Maximum likelihood trees were estimating using PhyML 3.0 (26), incorporating the best-fit amino acid substitution model, here JTT +G+F, with 1000 bootstrap replicates using the Montpellier Bioinformatics Platform (http://www.atgc-montpellier.fr/phyml/).

The avian HDV-like agent was subsequently interrogated for conserved features of human HDV. At the genomic level, GC content want calculated within Geneious using a sliding window of 10. To ascertain whether the circular genome folded into a classical unbranched rod sctructure we utilized a RNA folding algorithm implemented on the mfold webserver (27). The identification of the avian HDV-like agent ribozyme was performed in two phases. The TT2NE algorithm was first applied on the genomic and antigenomic ribozyme sequences of the reference HDV genome (GenBank accession X04451.1), and although the models inferred by TT2NE were inaccurate (13), their topologies were distinct and were used as references for screening. The free energy calculated for the HDV genomic and antigenomic ribozyme sequences were -47.35 kcal/mol and -36.70 kcal/mol, respectively. We screened incremental windows of 100 nt starting from first position of the antigenome of the avian HDV-like agent, also including the complementary genomic sequences, with the TT2NE algorithm to locate the potential ribozyme sequences. The incremental size was 50 nt from positions 1 to 500 of the genome sequence and then 10 nt from position 500. All inferred secondary structures that had free energy values similar to the reference models were further evaluated using PseudoViewer (28). Coiled-coil structures of the ORF were identified using Multicoil (29).

## Acknowledgements

We are grateful to members of the Centre for Integrative Ecology at Deakin University for sample collection and Melbourne Water for logistic support. This work was supported by an ARC Discovery Grant (DP160102146). ECH is supported by an ARC Australian Laureate Fellowship (FL170100022) and the Melbourne WHO Collaborating Centre for Reference and Research on Influenza is funded by the Australian Department of Health.

